# An endonuclease V protects *Bacillus subtilis* against a deaminase toxin and a uracil-containing phage

**DOI:** 10.64898/2026.09.29.755374

**Authors:** Adam J. Oliver, Stephan Joseph, Amit Singh, Ali Riazuddin, Marcus Lopez, Yang Gao, David Bates, Marcos H. de Moraes

## Abstract

Bacteria deploy polymorphic toxins to inhibit competitors and rely on cognate immunity proteins to prevent self-intoxication. Immunity determinants act by binding and neutralizing incoming toxins, and in the rare cases where toxin-induced damage is also reversed, that activity resides within the immunity protein itself. Whether a conflict system can instead dedicate a separate gene to repairing the damage is unknown. Here we report that an operon in *Bacillus subtilis* encodes the toxin YwqJ, its cognate immunity protein YwqK, and EndoV^Bs^ an endonuclease that incises DNA at the lesion the toxin creates. YwqJ is a single-stranded DNA cytidine deaminase translocated by the type VIIb secretion system (T7SSb) to mediate contact-dependent interbacterial antagonism. Unexpectedly, EndoV^Bs^ exhibits a preference for uracil over the canonical hypoxanthine, processing genomic uracil in cells and protecting against YwqJ when immunity is absent. Because EndoV^Bs^ acts on the chemical lesion rather than the effector, its protection is indifferent to the source of deamination. EndoV^Bs^ accordingly restricts replication of PBS1, a bacteriophage whose genome naturally substitutes uracil for thymine. These findings establish DNA repair as a novel layer of defense within toxin–immunity loci and reveal a DNA repair mechanism that unifies interbacterial antagonism and antiviral immunity.

## Introduction

Bacteria employ contact-dependent secretion systems to deliver polymorphic toxins into rival cells during interbacterial competition (1–3). In Gram-positive bacteria, the type VIIb secretion system (T7SSb) translocates diverse antibacterial effectors, typically encoded in tandem with cognate immunity factors that protect against self-intoxication (4, 5). While characterized immunity mechanisms rely on dedicated immunity proteins that neutralize the toxin (6–8), secondary protection mechanisms that resolve the resulting damage remain largely unexplored.

While polymorphic deaminase toxins have attracted significant attention(9–11), the substrates and base specificities of many predicted families remain unknown. In *B. subtilis*, the T7SSb-associated toxin YwqJ is predicted bioinformatically to contain a deaminase domain, but its catalytic activity has not been experimentally demonstrated. Adding to this ambiguity, the *ywqJK* operon encodes a third protein, YwqL, an endonuclease V family member (EndoV^Bs^). In organisms such as Escherichia coli and Thermotoga maritima, endonuclease V canonically initiates the repair of deaminated purines by cleaving DNA containing hypoxanthine. However, genetic studies in B. subtilis have linked EndoV^Bs^ to survival following exposure to chemical deaminating agents. Why a putative deaminase effector is genetically coupled to an endonuclease V, and whether EndoV^Bs^ acts to mitigate damage inflicted by its operon partner, remains unknown.

Here we define the activity of YwqJ and the function of its associated endonuclease. YwqJ is a cytidine deaminase that acts on single-stranded DNA, and its antagonism of *B. subtilis* requires both an intact T7SSb and an intact active site. EndoV^Bs^ cleaves uracil-containing DNA in preference to hypoxanthine and acts on genomic uracil in vivo. Cells lacking *ywqL* are sensitized to YwqJ and to the uracil-containing phage PBS1. These findings reveal that a repair enzyme acts alongside a dedicated immunity protein, extending protection against deaminase toxins beyond toxin neutralization.

## Results

### An uncharacterized deaminase among the *B. subtilis* LXG toxins

*Bacillus subtilis* strains encode an arsenal of polymorphic toxins containing an amino-terminal LXG trafficking domain, which are translocated into neighboring cells by the type VIIb secretion system (T7SSb) under biofilm-inducing conditions(12, 13). The carboxy-terminal domains of several LXG toxins act as nucleases, including YeeF, YobL, YokI, YqcG, and YxiD(12). However, the biochemical activity of YwqJ has remained undefined despite its demonstrated role in antagonism. The *ywq* locus exhibits the canonical architecture of a T7SSb substrate operon, containing the genes *ywqH* and *ywqI* encoding for small α-helical hairpin proteins, the toxin gene *ywqJ*, and the cognate immunity determinant *ywqK* (Fig. 1A). YwqJ comprises an amino-terminal LXG trafficking domain, a central PT-TG module, and a carboxy-terminal domain annotated as a putative deaminase (PF14431) (Fig. 1A). Immediately downstream of *ywqK*, and in the same transcriptional orientation, lies *ywqL*, encoding a member of the endonuclease V family (herein EndoV^Bs^).

**Figure 1.**
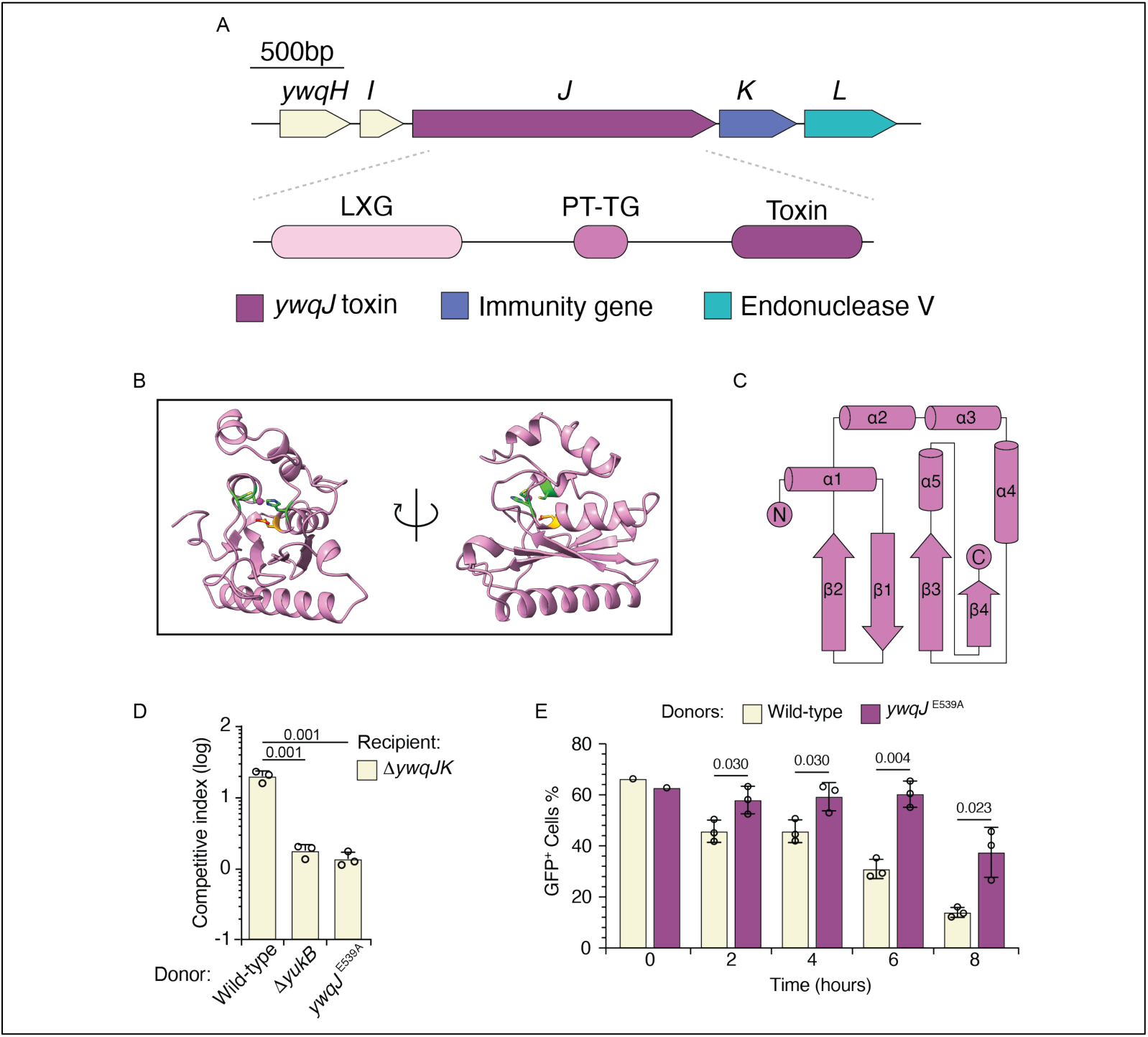
YwqJ is a T7SSb-delivered toxin with a predicted deaminase active site. **A**, Organization of the *ywqHIJKL* locus and domain architecture of YwqJ, showing the LXG trafficking domain, the pre-toxin PT-TG module and the C-terminal toxin domain. Scale bar, 500 bp. **B**, AlphaFold3 model of the YwqJ toxin domain, shown in two orientations related by a 180° rotation. The zinc-coordinating residues H537, C581 and C583 and the catalytic glutamate E539 surround a modeled zinc ion (sphere). **C**, Topology of the toxin domain, showing the five-stranded β-sheet (β1-β4) and flanking helices (α1-α5). **D**, Competitive index of a Δ*ywqJK* recipient co-cultured with wild-type, Δ*yukB* or *ywqJ*^E539A^ donor strains. Bars show the mean of three biological replicates with individual values overlaid; error bars, s.d. **E**, Proportion of GFP^+^ Δ*ywqJK* recipient cells during co-culture with wild-type or *ywqJ*^E539A^ donors, measured by flow cytometry over 8 h. n = 3 biological replicates; error bars, s.d. Statistical significance was assessed by ordinary one-way ANOVA with Tukey’s multiple comparisons test (**D**) or an unpaired two-tailed Student’s *t*-test (**E**).

To examine the uncharacterized toxin domain, we generated a structural model using AlphaFold3 (14) (Fig. 1B, C, Supplementary Fig. 1). The model was predicted with high confidence and adopted a core deaminase fold consisting of a five-stranded β-sheet flanked by buttressing α-helices(15, 9) (Fig. 1C). Conserved zinc-coordinating residues (H537, C581, and C583) together with a predicted catalytic glutamate (E539) clustered within a single pocket coordinating a modeled zinc ion(16) (Fig. 1B). Although YwqJ displays two additional central helices and an extended N-terminal helix relative to other characterized bacterial deaminase toxin folds, this active-site architecture suggested that the enzyme functions as a zinc-dependent deaminase with E539 serving as the catalytic proton donor.

We asked whether competitive inhibition mediated by YwqJ requires both an intact secretion apparatus and the predicted deaminase active site. We constructed a sensitized recipient strain lacking the toxin–immunity pair (Δ*ywqJK*) carrying a constitutive GFP reporter, and competed it against wild-type, a secretion-deficient mutant lacking the core T7SSb component YukB(13) (Δ*yukB*), or a donor strain carrying a catalytic substitution in the toxin active site (*ywqJ*^E539A^). The wild-type donor outcompeted the recipient by several orders of magnitude (Fig. 1D). This competitive advantage was completely lost in both the Δ*yukB* and *ywqJ*^E539A^ donors, with competitive indices that were indistinguishable from one another (Fig. 1D).

Intoxication kinetics tracked by flow cytometry confirmed this requirement. The proportion of GFP+ recipient cells declined from approximately 65% to 14% over 8 h in co-culture with wild-type donor, whereas recipients survived in the presence of the *ywqJ*^E539A^ strain (Fig. 1E). Interbacterial antagonism by YwqJ therefore occurs rapidly upon cell contact and strictly depends on both a functional T7SSb apparatus and the predicted catalytic residue.

### YwqJ mediates T7SSb-dependent antagonism

We sought to establish whether antagonism requires both the secretion apparatus and the predicted deaminase active site. We generated a sensitized recipient strain lacking the toxin-immunity pair, Δ*ywqJK*, and labeled it with constitutively expressed GFP, then competed this recipient against a wild-type donor, a Δ*yukB* donor lacking an essential component of the T7SSb apparatus, and a donor carrying a single substitution of the predicted catalytic residue (*ywqJ*^E539A^). The wild-type donor exhibited a marked competitive advantage against the recipient (Fig. 1D). This advantage was not observed for the Δ*yukB* or the *ywqJ*^E539A^ donor, and the competitive indices of these two strains did not differ significantly from one another (Fig. 1D). These data indicate that YwqJ-dependent antagonism requires a functional T7SSb and an intact deaminase active site.

We next examined the kinetics of intoxication by tracking the recipient population using flow cytometry. In co-culture with the wild-type donor, the proportion of GFP^+^ recipient cells declined from approximately 65% to 14% over 8 h (Fig. 1E). By contrast, in co-culture with the *ywqJ*^E539A^ donor, the proportion of GFP^+^ cells remained significantly higher at each time point, with the difference apparent by 2 h (Fig. 1E). Together, these findings indicate that YwqJ-dependent killing occurs rapidly following cell contact and requires the predicted catalytic residue.

### YwqJ is a cytidine deaminase that acts on ssDNA

Neither the structural fold nor the active-site arrangement of YwqJ identifies its target. The majority of the bacterial DNA deaminase toxin families (BaDTFs) characterized to date target DNA(10, 11, 17, 18), and we therefore evaluated DNA substrates first. Guided by the structural model, we separated the toxin domain (YwqJ_tox_) from the LXG and PT-TG trafficking modules, co-expressed YwqJ_tox_ with the immunity protein YwqK in *E. coli* and separated the two proteins by on-column denaturation and refolding.

We measured the catalytic activity of purified YwqJ_tox_ on a 37-nucleotide single-stranded DNA (ssDNA) substrate containing multiple cytidines (A15-ACTCGCC-A15). YwqJ_tox_ deaminated this substrate efficiently, with activity detected at enzyme concentrations as low as 40 nM, whereas the catalytically inactive variant YwqJ_tox_(E539A) yielded no detectable product (Fig. 2A). Multiple product bands were generated in the ssDNA reaction, indicating that YwqJ_tox_ deaminates more than one cytidine within the substrate. By contrast, activity on the corresponding double-stranded DNA (dsDNA) substrate was minimal (Fig. 2B). Control deaminases confirmed these preferences. SsdA_tox_, an established ssDNA cytidine deaminase, was active exclusively on the single-stranded substrate, whereas DddA_tox_ was active only on dsDNA (Fig. 2A, B).

**Figure 2.**
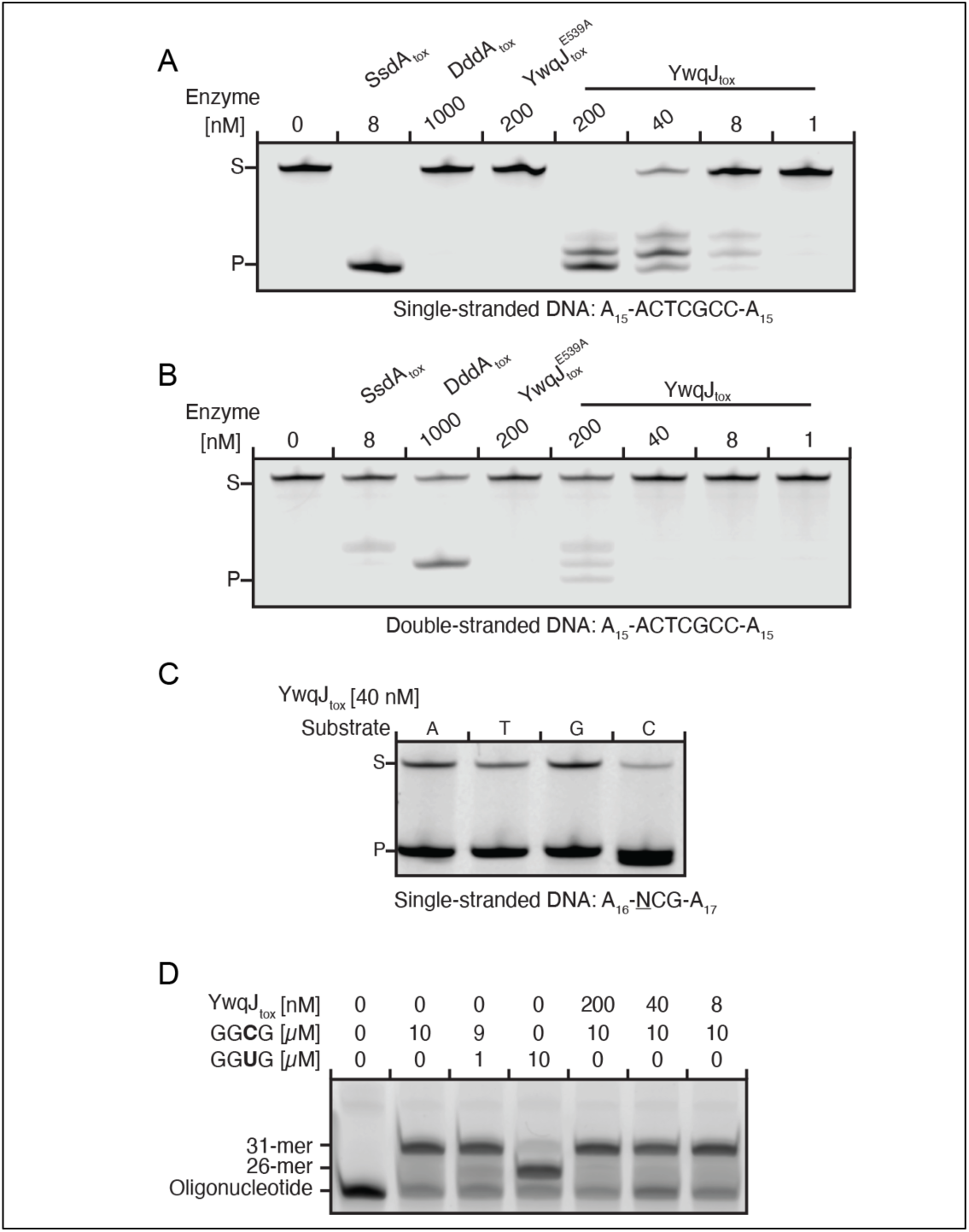
YwqJ is a cytidine deaminase that acts on single-stranded DNA. **A**, Deamination of a labeled single-stranded DNA substrate (A_15_-ACTCGCC-A_15_) by the indicated toxin domains at the concentrations shown. Reactions were treated with uracil-DNA glycosylase and NaOH before denaturing gel electrophoresis, such that the product (P) reports deaminated cytidine; S, substrate. SsdA_tox_ and DddA_tox_ are included as single- and double-stranded DNA-specific controls, respectively, and YwqJ_toxE539A_ as a catalytically inactive control. **B**, As in **A**, with the corresponding double-stranded substrate. **C**, Deamination by YwqJ_tox_ of single-stranded substrates carrying a single cytidine in each of the four 5′ flanking-base contexts (A_16_-NCG-A_17_); the flanking base is underlined. **D**, Poisoned primer extension on an RNA oligonucleotide containing the target cytidine within a GGCG motif. An otherwise identical oligonucleotide synthesized with uridine in place of the target cytidine (GGUG) models the deaminated product and arrests extension at 26 nucleotides, whereas unmodified template yields the full-length 31-nucleotide product.

Characterized DNA cytidine deaminases frequently operate with a preference for the nucleotide immediately 5′ of the target cytidine(19, 20). We asked whether YwqJ shares this requirement and measured its activity on ssDNA substrates containing a single cytidine in each of the four possible 5′-NC contexts (A12-NCG-A17). YwqJ_tox_ deaminated all four substrates with comparable efficiency (Fig. 2C).

These results indicate that, in contrast to most characterized BaDTF members, YwqJ does not require a specific 5′ flanking base. Because deaminases can act on RNA as well as DNA(21, 17), we next asked whether YwqJ deaminates RNA substrates. Using a poisoned primer extension assay, we tested YwqJ_tox_ against an RNA oligonucleotide containing a target cytidine within a GGCG motif (Fig. 2D). In the absence of an available positive-control RNA deaminase, we benchmarked detection using an identical oligonucleotide synthesized with uridine in place of the target cytidine (GGUG), which models a deaminated product and directs primer extension arrest at the 26-nucleotide position. While the synthetic GGUG control produced the expected 26-nucleotide product, the GGCG substrate incubated with YwqJarrest at the 26-nucleotide position. While the synthetic GGUG control produced the expected 26-nucleotide product, the GGCG substrate incubated with YwqJ_tox_ yielded exclusively the full-length 31-nucleotide product, indistinguishable from untreated RNA (Fig. 2D). These findings establish that YwqJ deaminates single-stranded DNA rather than RNA.

Previous screens of YwqJ-family deaminases did not detect activity on ssDNA or dsDNA. Two methodological differences may account for this discrepancy(11, 17). First, the enzymes examined previously share less than 22% sequence identity with YwqJ_tox_, raising the possibility that members of the family differ in substrate preference. Second, those studies relied on split-deaminase constructs in cell culture or on unpurified cell-free transcription-translation extracts. Poor heterologous expression, improper folding, or unoptimized reaction conditions in such formats can yield false negatives, which we avoided by assaying the purified toxin domain directly.

The identification of YwqJ as a ssDNA deaminase bears on the role of DNA deamination in interbacterial antagonism. Several members of the bacterial deaminase superfamily have been adapted for genome editing, and we have characterized SsdA as a ssDNA cytidine deaminase predicted to be secreted by the T6SS of *Pseudomonas syringae*(*22*). However, DddA of *B. cenocepacia*, a dsDNA deaminase(10), and the umbrella particle UmbC1 from *Streptomyces coelicolor*(*23*), both from the same BaDTF, were previously the only members of the superfamily whose physiological role as a secreted toxin had been directly demonstrated. Our finding that YwqJ acts as a potent ssDNA deaminase during contact-dependent competition indicates that ssDNA deamination is a physiologically relevant strategy in interbacterial antagonism. The divergence of the YwqJ structure from that of other BaDTF members is accompanied by a divergence in substrate preference, which suggests that the enzyme engages its substrate through contacts not present in the characterized members of the superfamily.

### EndoV^Bs^ preferentially cleaves uracil-containing DNA

*DNA Immediately downstream of the cognate immunity gene ywqK*, and in the same transcriptional orientation, lies *ywqL*, which encodes a member of the endonuclease V family (EndoV^Bs^). In previously characterized polymorphic toxin systems, immunity is mediated exclusively by binding proteins that neutralize the toxin. The presence of a co-operonic DNA repair endonuclease is without precedent, suggesting that YwqJ and EndoV^Bs^ act on the same lesion.

In bacteria, endonuclease V canonically initiates the repair of deaminated purines, specifically hypoxanthine(24). The enzyme hydrolyzes the second phosphodiester bond located 3′ of the lesion to generate a single-strand nick(25), after which DNA polymerase I excises the damaged strand and DNA ligase seals the break (24). In *E. coli* and *Thermotoga maritima*, hypoxanthine is the preferred physiological substrate, although purified enzymes can process other lesions *in vitro*(26–28) (29). Because YwqJ deaminates cytidine to generate uracil, we asked whether EndoV^Bs^ has evolved a preference for uracil over hypoxanthine.

We purified recombinant EndoV^Bs^ and compared its cleavage activity on matched oligonucleotide substrates containing either uracil or hypoxanthine at a single position (A15-N-A15). On ssDNA, EndoV^Bs^ cleaved both substrates, but activity was markedly higher on the uracil-containing template, remaining detectable down to 0.1 μM enzyme where hypoxanthine cleavage was lost (Fig. 3A). This preference was pronounced on dsDNA, where the U:A substrate was cleaved substantially more efficiently than the Hx:G substrate (Fig. 3B). This outcome was unexpected, because U:A forms a stable Watson–Crick base pair, whereas Hx:G is an unstable mismatch. By contrast, parallel assays with purified *E. coli* Nfi confirmed its canonical preference for hypoxanthine over uracil (Supplementary Fig. 2). Thus, EndoV^Bs^ exhibits a different substrate specificity relative to canonical endonuclease V, favoring uracil over hypoxanthine.

**Figure 3.**
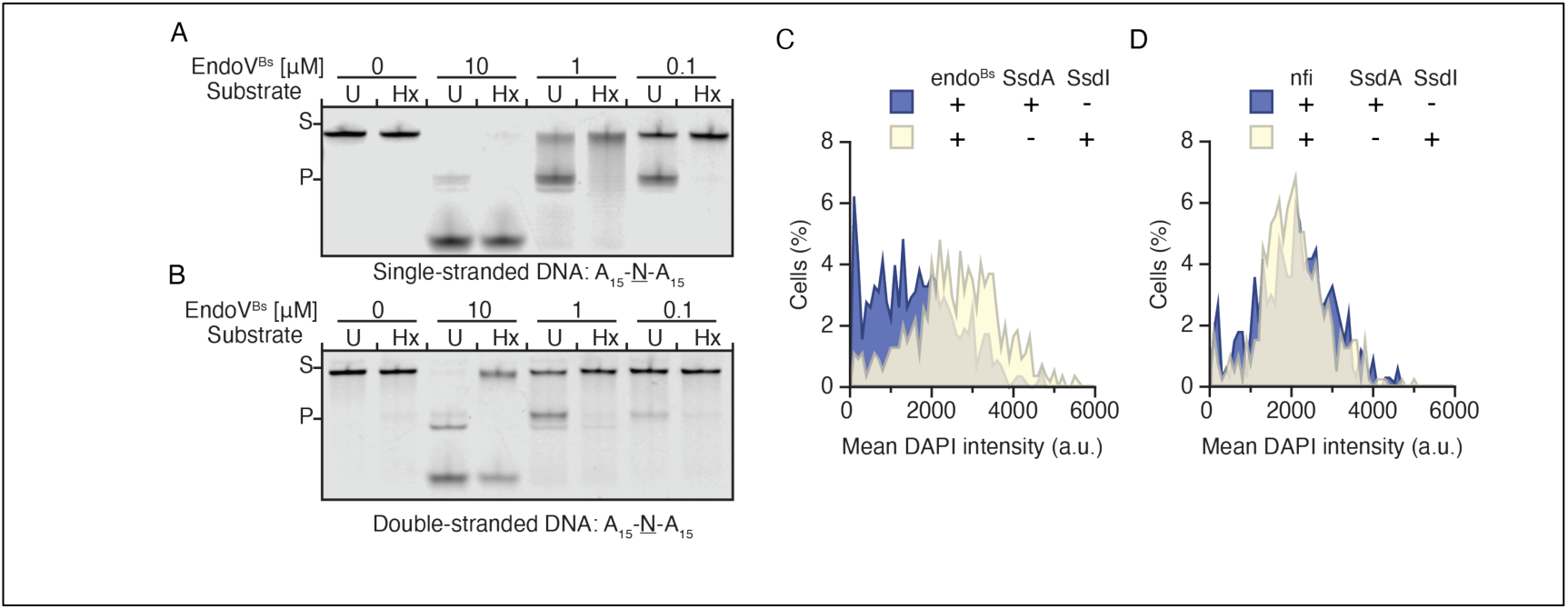
EndoV^Bs^ cleaves uracil-containing DNA in vitro and processes genomic uracil in cells. **A**, Cleavage of labeled single-stranded DNA substrates containing a single uracil (U) or hypoxanthine (Hx) (A_15_-N-A_15_) by EndoV^Bs^ at the indicated concentrations. S, substrate; P, product. **B**, As in **A**, with double-stranded substrates in which the modified base is paired with adenine (dU:A) or guanine (dI:G). **C**, Distribution of mean DAPI fluorescence intensity in single *E. coli* Δ*ung* Δ*nfi* cells expressing EndoV^Bs^ together with SsdA (blue) or with SsdA neutralized by its cognate immunity protein SsdI (cream). **D**, as in **C**, with Nfi expressed in place of EndoV^Bs^.

### EndoV^Bs^ acts on genomic uracil *in vivo*

Cleavage in vitro does not establish that an enzyme processes that lesion in cells. For example, Nfi cleaves uracil *in vitro* but does not act on uracil in *in vivo*(*28*). To determine whether EndoV^Bs^ acts on genomic uracil in living cells, we established a defined heterologous test system in E. coli. Because ectopic overexpression of YwqJ causes non-specific toxicity, we supplied genomic uracil using SsdA, a potent ssDNA cytidine deaminase that functions efficiently in *E. coli*(22). We expressed SsdA alongside either EndoV^Bs^ or Nfi in an *E. coli* Δ*ung* Δ*nfi* background. Deleting *ung* prevents canonical base excision repair of uracil, ensuring that endonucleolytic cleavage introduces unsealed single-strand nicks that lead to chromosome degradation. Deleting *nfi* removes the endogenous endonuclease V, providing an otherwise null background.

We quantified nucleoid integrity in individual cells using DAPI fluorescence microscopy, comparing cells expressing active SsdA to controls in which SsdA was neutralized by its cognate immunity protein SsdI. In cells expressing EndoV^Bs^, active cytidine deamination generated a distinct subpopulation of cells (approximately 6%) with near-zero mean DAPI intensity, indicative of chromosome fragmentation and nucleoid loss (Fig. 3C). This degraded subpopulation was absent in the SsdI-neutralized control. By contrast, when Nfi was substituted for EndoV^Bs^, no degraded subpopulation emerged, and the DAPI intensity distributions between the active and neutralized conditions remained superimposable (Fig. 3D). These findings demonstrate that EndoV^Bs^, but not canonical Nfi, recognize and process genomic uracil in living cells.

### EndoV^Bs^ protects *B. subtilis* against YwqJ intoxication

Having established that EndoV^Bs^ cleaves uracil-containing DNA in vitro and in cells, we assessed whether it protects *B. subtilis* against cytidine deamination inflicted by YwqJ. In competition assays between immunity-intact strains, deletion of *endoV*^Bs^ alone conferred no competitive disadvantage. A Δ*endoV*^Bs^ recipient displayed wild-type fitness when co-cultured with either a wild-type donor or a catalytically inactive *ywqJ*^E539A^ donor (Fig. 4A). This outcome is expected, because the recipient retains the cognate immunity gene *ywqK* and is protected from delivered YwqJ.

**Figure 4.**
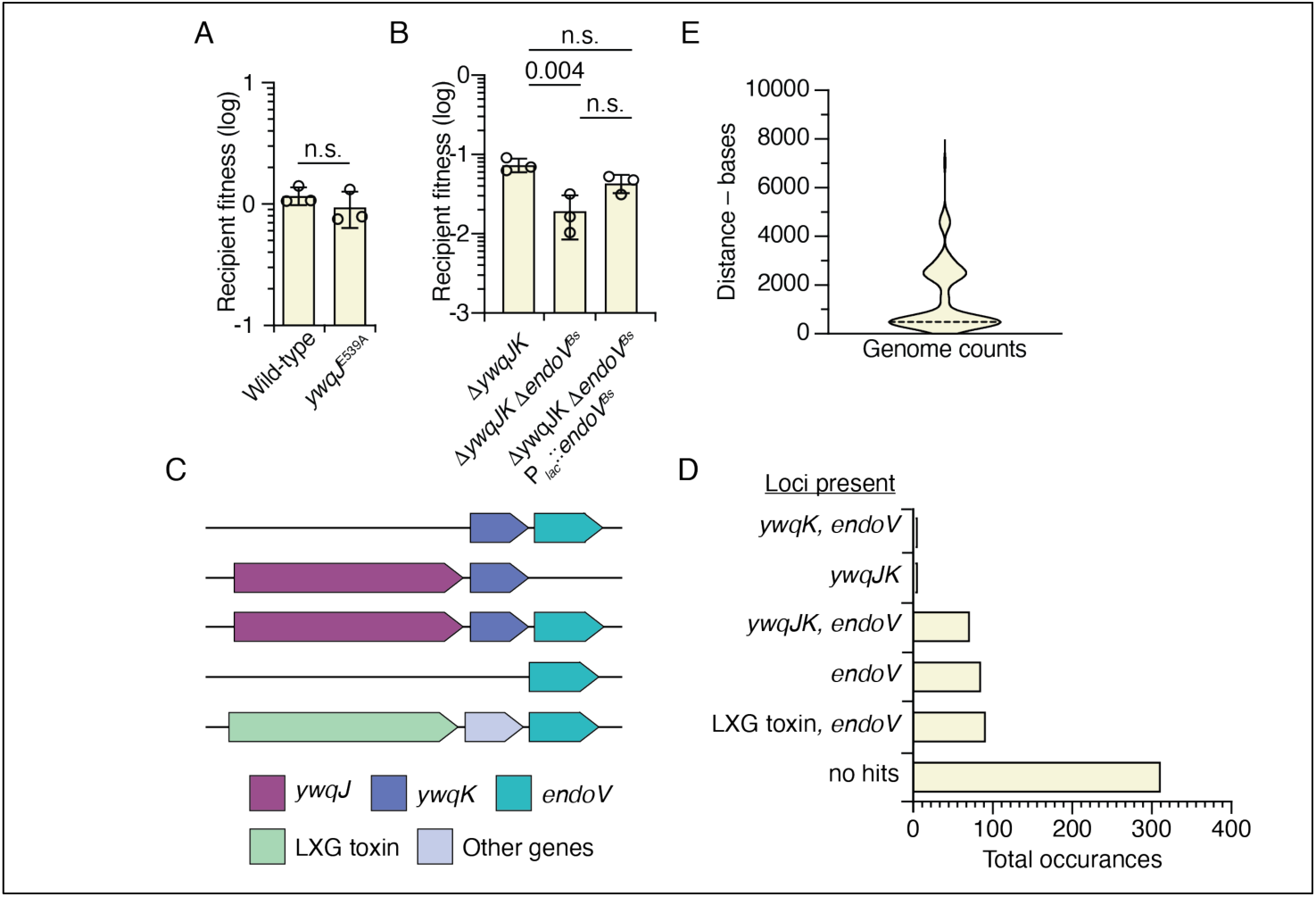
EndoV^Bs^ protects *B. subtilis* against YwqJ intoxication and is genetically linked to *ywq* toxin loci. **A**, Fitness of a Δ*endoV*^Bs^ recipient after co-culture with a wild-type or *ywqJ*^E539A^ donor. **B**, Fitness of Δ*ywqJK*, Δ*ywqJK* Δ*endoV*^Bs^ and complemented Δ*ywqJK* Δ*endoV*^Bs^ P_lac_::*endoV*^Bs^ recipients after co-culture with a wild-type donor. **C**, Gene neighbourhood architectures containing *ywqJ*, *ywqK*, *endoV* or unrelated LXG toxin genes, identified across 449 complete *B. subtilis* genomes using Pandoomain. **D**, Occurrences of each architecture; genomes lacking all query genes are shown as no hits. **E**, Intergenic distance between co-oriented *endoV* and LXG toxin genes; dashed line, median. In **A** and **B**, bars show the mean of three biological replicates with individual values overlaid; error bars, s.d. Statistical significance was assessed by ordinary one-way ANOVA with Tukey’s multiple comparisons test. n.s., not significant.

To unmask any contribution from the endonuclease, we removed the immunity determinant from the recipient. We competed a wild-type donor against sensitized recipients lacking immunity (Δ*ywqJK*), lacking both immunity and the repair enzyme (Δ*ywqJK* Δ*endoV*^Bs^), or complemented with the endonuclease in trans (Δ*ywqJK* Δ*endoV*^Bs^ Plac::*endoV*^Bs^). Recipient survival was impaired in all three competitions. However, loss of *endoV*^Bs^ significantly exacerbated this killing relative to the Δ*ywqJK* control (Fig. 4B). Complementation with *endoV*^Bs^ expressed from an inducible promoter restored recipient fitness to that of the Δ*ywqJK* strain (Fig. 4B). Thus, EndoV^Bs^ provides a functional second layer of defense in B. subtilis, conferring resilience against YwqJ-mediated cytidine deamination when immunity binding is absent or overwhelmed.

Cytidine deamination lesions generated by YwqJ in recipient cells escaped direct physical detection. We did not detect elevated genomic uracil or C-to-T transition mutations in bulk genomic DNA extracted from intoxicated recipients, even in a sensitized Δ*ung* Δ*endoV*^Bs^ background lacking both characterized uracil-processing pathways. This outcome is consistent with the biochemical properties of the toxin. YwqJ acts strictly on single-stranded DNA, which is transiently exposed during replication and transcription, representing a minute fraction of the chromosome at any moment. As a result, population-wide lesion density remains below the threshold of bulk sequencing methods, while remaining lethal to individual intoxicated cells. The evidence that YwqJ deaminates DNA in recipients is therefore genetic: contact-dependent killing requires the catalytic glutamate E539 (Fig. 1D), and loss of the uracil-directed endonuclease EndoV^Bs^ specifically sensitizes recipients to the toxin (Fig. 4B).

### Endonuclease V genes are associated with *ywq* toxin loci

Because EndoV^Bs^ mitigates damage from YwqJ-mediated cytidine deamination, we asked whether this functional pairing is reflected in genome organization. Using the Pandoomain annotation pipeline(30), we systematically surveyed the co-localization of genes encoding YwqJ, cognate immunity YwqK, other LXG-family toxins, and EndoV^Bs^ across all 449 complete *B. subtilis* genomes available on the National Center for Biotechnology Information (NCBI) database.

This analysis identified five distinct gene neighborhood architectures with contrasting abundances (Fig. 4C, D). These comprised 93 occurrences of solitary *endoV*, 87 of *endoV* paired with an unrelated LXG toxin, 43 of the canonical *ywqHIJKL* operon, 3 of *ywqJK* lacking the endonuclease, and 1 of *endoV* coupled solely to *ywqK*. The remaining 313 genomes lacked all query genes. Where *endoV* and an LXG toxin gene co-occur, the two genes are consistently co-oriented and separated by a median distance of approximately 487 base pairs (Fig. 4E).

These architectures reveal several features of the locus that bear on its physiological function. The *ywq* locus is dynamic, with toxin domains gained and lost while the amino-terminal LXG trafficking module remains conserved. Notably, *endoV* is absent from approximately 70% of the genomes examined. Such a sporadic distribution is unexpected for a general housekeeping repair enzyme and instead points to a specialized role in interbacterial antagonism. Wherever *ywqJK* is present, *endoV* is almost invariably situated immediately downstream, consistent with the protective phenotype observed in competition. The converse does not hold. Endonuclease V occurs alone, or adjacent to loci encoding alternative LXG toxins, as frequently as it occurs with *ywqJK*. Among these alternative loci, those with recognizable annotations encode ribonucleases or classical endonucleases (Supplementary Table 1). However, many of the LXG effectors paired with *endoV* returned no functional domain matches, indicating that they share low sequence homology with existing database alignments or represent entirely new toxin families. These bioinformatic patterns indicate that *endoV*^Bs^ is tightly linked to *ywqJ* but can be retained independently, suggesting that the enzyme may also confer generic protection against cytidine deaminases delivered by competing strains.

### EndoV^Bs^ restricts replication of the uracil-containing phage PBS1

Because EndoV^Bs^ recognizes the lesion rather than the toxin that produced it, DNA bearing uracil should serve as a substrate regardless of origin. *B. subtilis* is susceptible to infection by PBS1, a bacteriophage whose genome naturally substitutes uracil for thymine(31). This substitution renders PBS1 DNA refractory to many restriction endonucleases that act on conventional DNA, and is presumed to contribute to evasion of host defense(32). To preserve this modification during infection, PBS1 and its clear-plaque derivative PBS2 express a potent uracil-DNA glycosylase inhibitor (Ugi) that binds and inactivates host Ung, preventing the enzymatic excision of uracil from viral DNA(33). Because EndoV^Bs^ cleaves double-stranded DNA uracil independently of Ung, we asked whether it restricts PBS1 infection.

To evaluate host susceptibility across distinct physiological states, we challenged cultures with PBS1 at multiplicities of infection (MOI) of 0.1 and 1 in either the exponential phase or stationary phase. For exponential-phase assays, phage was added directly to actively growing cultures (2 to 3 h of growth). Under these conditions, wild-type cultures exhibited rapid lysis at both high viral challenge (MOI 1) and at low multiplicity (MOI 0.1) (Fig. 5A, B). For stationary-phase assays, cells from 24 h stationary cultures were inoculated into fresh LB broth at the point of infection. Because stationary cells must exit quiescence before resuming active division, the onset of phage replication and subsequent lysis was substantially delayed, resulting in prolonged maintenance of optical density for the wild-type host (Fig. 5C, D).

**Figure 5.**
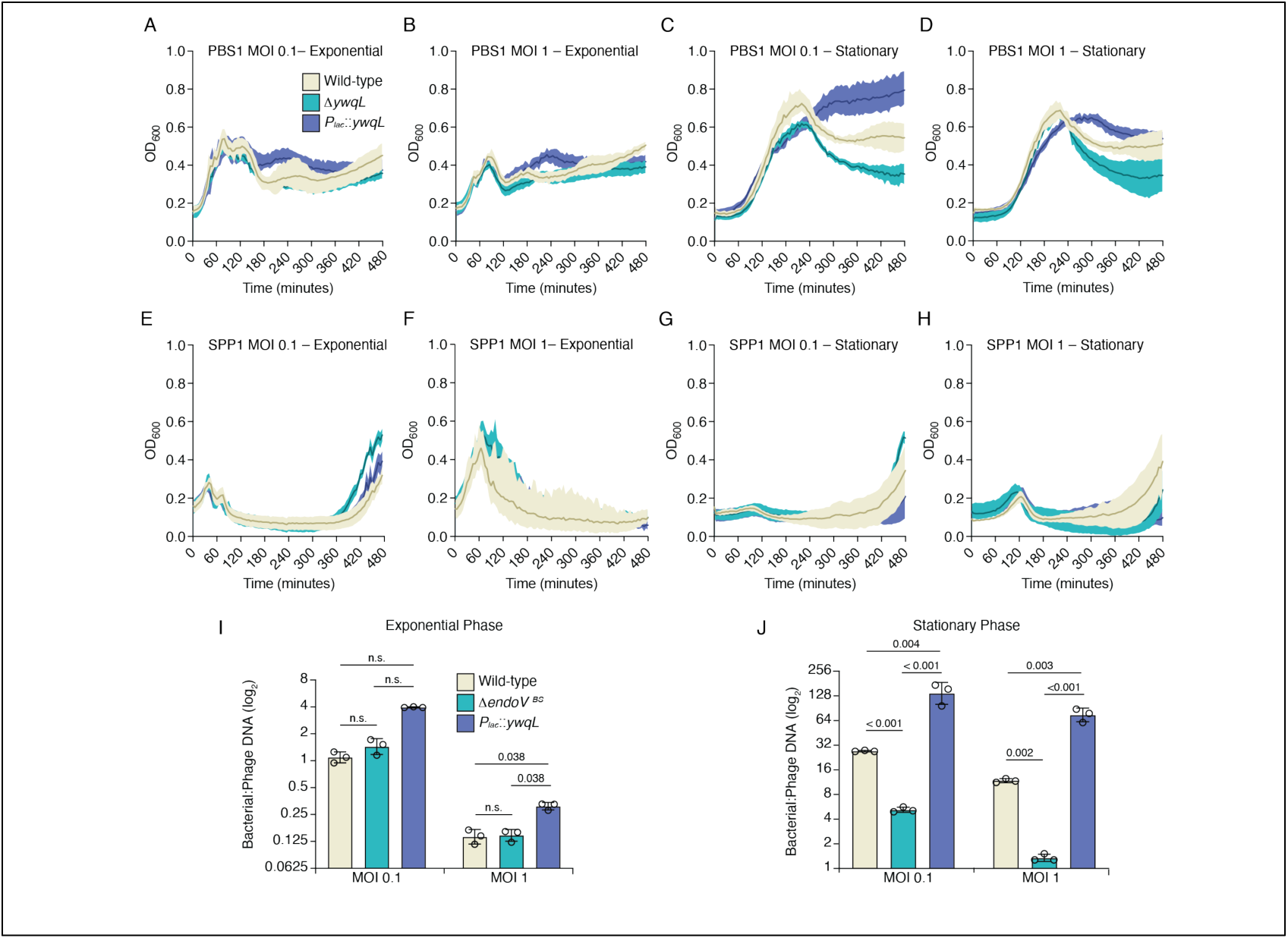
EndoV^Bs^ restricts replication of the uracil-containing phage PBS1. **A**–**D**, Growth of wild-type, Δ*endoV*^Bs^ and complemented (P_lac_::*endoV*^Bs^) *B. subtilis* challenged with PBS1 at a multiplicity of infection (MOI) of 0.1 or 1, either in exponential phase (**A**, **B**) or after transfer from stationary phase (**C**, **D**). **E**–**H**, as in **A**–**D**, with the thymine-containing phage SPP1. Lines show the mean of three biological replicates; shaded areas, s.d. **I**, **J**, Ratio of bacterial to phage DNA determined by quantitative PCR for cultures infected in exponential phase (**I**) or after transfer from stationary phase (**J**), sampled 30 min after phage addition. Bars show the mean of three biological replicates with individual values overlaid; error bars, s.d. Statistical significance was assessed by Brown-Forsythe ANOVA with Dunnett’s T3 multiple comparisons test. n.s., not significant.

Evaluating the endonuclease-deficient mutant (Δ*endoV*^Bs^) and the complemented strain (Δ*endoV*^Bs^ Plac::*endoV*^Bs^) against the wild-type baseline revealed that EndoV^Bs^ affects the severity of lysis and subsequent recovery rather than the timing of infection. When stationary-phase cells were transferred into fresh medium and challenged with PBS1, all strains began lysing at approximately 210 minutes after infection (Fig. 5C, D). However, the magnitude of lysis and the capacity for regrowth differed substantially across the three strains. While wild-type cultures exhibited limited lysis before arresting culture decline and resuming growth, cultures of the Δ*endoV*^Bs^ mutant suffered far more pronounced lysis and collapsed without recovering (Fig. 5C, D). Ectopic expression of the endonuclease in trans completely rescued this phenotype (Fig. 5C, D).

Comparing growth curves by calculating the area under the curve (A.U.C.) confirmed that the mutant’s pronounced lysis and failure to recover were statistically significant at both MOI 0.1 and MOI 1 (Supplementary Fig. 4C, D). Complementation restored total growth yields to wild-type levels at MOI 1 and elevated recovery significantly above the wild-type baseline at MOI 0.1, demonstrating a dose-dependent protective effect (Supplementary Fig. 4C, D). In exponential phase, all strains likewise initiated lysis simultaneously. Under high viral challenge (MOI 1), overall lysis trajectories and A.U.C. differences between wild type and mutant were not statistically significant (Fig. 5B, Supplementary Fig. 4B). However, under lower viral pressure (MOI 0.1), quantifying the A.U.C. revealed a modest but statistically significant defect in mutant recovery relative to wild-type that was restored by complementation (Supplementary Fig. 4A). Thus, EndoV^Bs^ does not alter the onset of phage-induced lysis, but restricts the depth of lysis and promotes host recovery, a protective effect that is detectable at low viral challenge in exponential phase and becomes pronounced following stationary-phase transfer.

To determine whether the observed protection depends specifically on uracil or represents an unspecific host response that broadly inhibits phage replication, we asked whether protection extends to SPP1, a *B. subtilis* phage with a conventional thymine-containing genome(34). We challenged wild-type, Δ*endoV*^Bs^, and complemented strains with SPP1 under the same conditions, examining both multiplicities of infection during exponential growth and following stationary-phase transfer (Fig. 5E–H). In contrast to the divergence observed with PBS1, all three strains displayed indistinguishable lysis kinetics across every condition tested. Comparing total growth yields by area under the curve confirmed that the presence or absence of EndoV^Bs^ had no measurable effect on culture density during SPP1 infection, with no statistically significant differences observed between any of the strains (Supplementary Fig. 4E–H). These results establish that the protection conferred by EndoVobserved between any of the strains (Supplementary Fig. 4E–H). These results establish that the protection conferred by ^Bs^ is not an unspecific antiviral state, but strictly requires the presence of uracil in the viral genome.

To determine whether the recovery of cell density reflects direct inhibition of viral replication, we quantified the relative abundance of bacterial to phage DNA by quantitative PCR (qPCR). Following transfer from stationary phase, the bacterial-to-phage DNA ratio differed dramatically across strains, mirroring the divergence in growth recovery (Fig. 5J). In wild-type cells, host DNA was maintained at a high ratio relative to viral DNA. In the Δ*endoV*^Bs^ mutant, this ratio dropped at both low and high multiplicities of infection, demonstrating that viral genomes accumulate to much higher levels in the absence of the endonuclease (Fig. 5J). Ectopic expression of EndoV^Bs^ (Plac::*ywqL*) elevated the bacterial-to-phage DNA ratio significantly beyond wild-type levels at both viral doses, confirming a dose-dependent restriction of viral replication (Fig. 5J). By contrast, in exponential-phase cultures, the bacterial-to-phage DNA ratio was substantially lower across all strains, and the difference between wild type and the Δ*endoV*^Bs^ mutant was not statistically significant (Fig. 5I). Thus, EndoV^Bs^ protects *B. subtilis* by directly suppressing the accumulation of phage PBS1 genomes, an effect that operates with high efficiency following stationary-phase transfer.

Several features of the stationary-phase physiology likely account for the pronounced restriction observed after transfer from the stationary phase. Expression of the *ywq* operon increases approximately three-fold as cultures enter stationary phase (Supplementary Fig. 2), consistent with reports that transcription of the operon is reduced in a quorum-sensing mutant(35). Upon dilution into fresh medium, cells pass through a lag phase, a distinct physiological state in which the transcriptional program is extensively remodeled before exponential growth(36). Incoming viral genomes thus enter a host in which EndoV^Bs is already present but which has not yet resumed rapid proliferation. In exponentially growing cells the reverse holds. EndoV^Bs^ levels are lower, and viral replication is rapid, such that cleavage of uracil-containing phage DNA is outpaced. We therefore propose that restriction reflects the ratio of EndoV^Bs^ activity to the rate of viral genome replication, which is most favorable to the host during outgrowth from the stationary phase.

### Loss of EndoV^Bs^ is costly during PBS1 infection

Monoculture growth assays demonstrate individual host recovery, but they may underreport the evolutionary consequences of viral exposure in mixed populations. To determine whether EndoV^Bs^ confers a competitive fitness advantage during infection, we competed a Δ*endoV*^Bs^ mutant against wild type in co-cultures challenged with PBS1. In stationary phase, the competitive index of the mutant declined progressively with increasing viral pressure, revealing a pronounced, dose-dependent fitness defect after 2 h of infection that was maintained after 4 h (Fig. 6C). In exponential phase, competitive indices trended downward with viral challenge, but these differences were not statistically significant (Fig. 6A). Loss of EndoV^Bs^ therefore imposes an acute competitive fitness cost during PBS1 challenge, mirroring the phase-dependent protection observed in monoculture.

**Figure 6.**
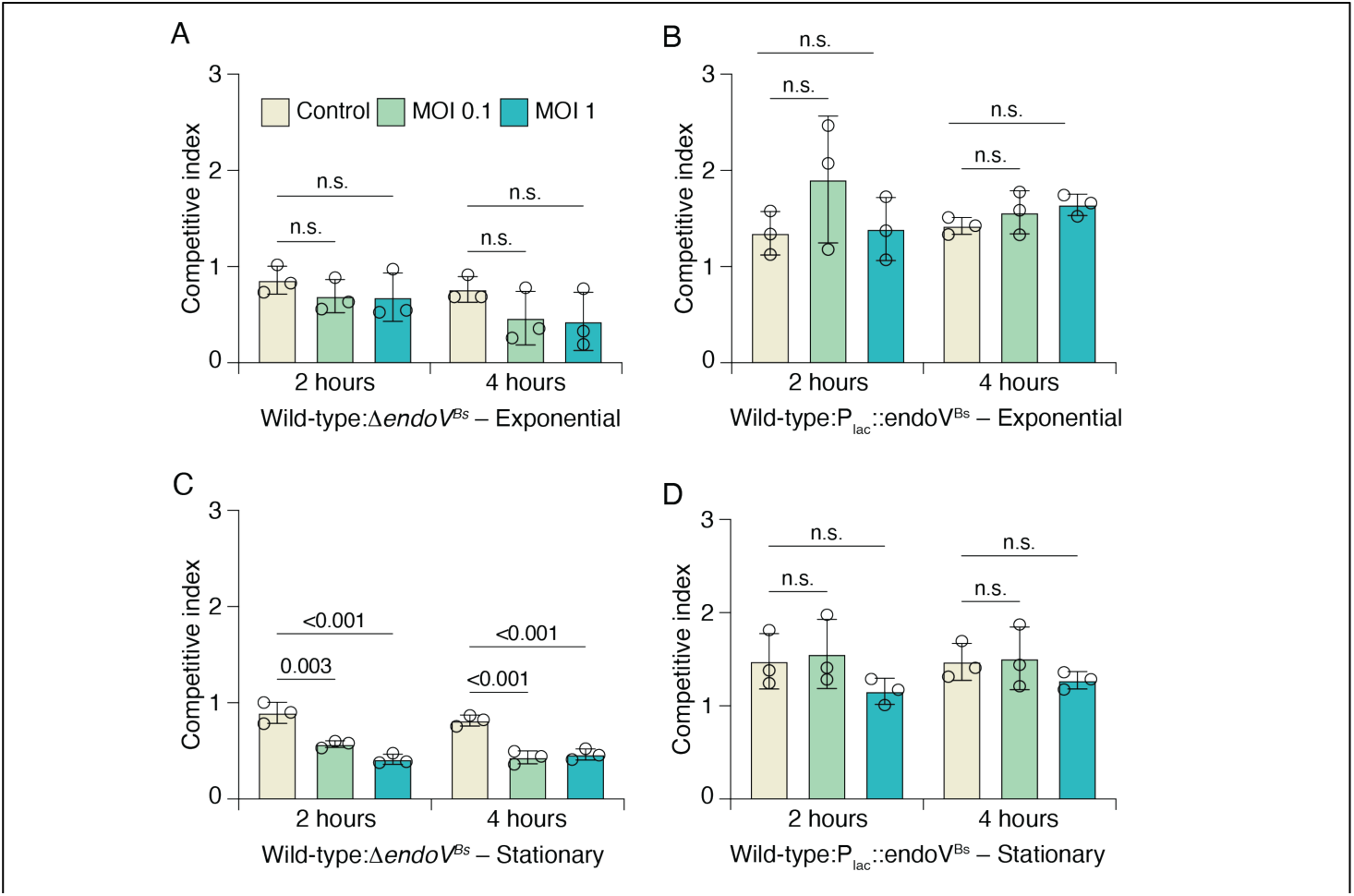
EndoV^Bs^ confers a competitive advantage during phage infection. **A**, **B**, Competitive index of a Δ*endoV*^Bs^ mutant (**A**) or a complemented P_lac_::*endoV*^Bs^ strain (**B**) co-cultured with wild type in exponential phase, either uninfected (control) or challenged with PBS1 at an MOI of 0.1 or 1, after 2 and 4 h. **C**, **D**, As in **A** and **B**, following transfer from stationary phase. Bars show the mean of three biological replicates with individual values overlaid; error bars, s.d. Statistical significance was assessed by ordinary one-way ANOVA with Tukey’s multiple comparisons test. n.s., not significant.

We also competed a complemented strain expressing *endoV*^Bs^ in trans against wild type. The competitive index of the complemented strain remained slightly above unity across both growth phases, unaffected by phage addition or viral multiplicity (Fig. 6B, D). Competitions performed with reciprocal antibiotic resistance markers confirmed that this slight deviation was already present in uninfected controls, reflecting a neutral marker bias rather than enhanced antiviral defense. Overexpressing EndoV^Bs^ therefore does not provide an additional fitness benefit in direct competition. Native levels of EndoV^Bs^ are sufficient to preserve competitive fitness during PBS1 infection, extending the protective role of the enzyme from competitor-derived cytidine deamination to viral challenge.

Our genomic survey revealed that *endoV* is absent from approximately 70% of sequenced *B. subtilis* isolates (Fig. 4D). In natural soil and plant-associated environments where strains harboring or lacking the *ywq* operon frequently co-occur, exposure to uracil-containing bacteriophages would impose strong asymmetric selective pressure. By selectively depleting strains that lack the endonuclease while sparing those that encode it, uracil-containing phages may directly dictate intraspecies population dynamics and drive the maintenance of this non-operon in polymicrobial communities.

## Discussion

The *ywq* locus encodes a DNA deaminase toxin together with an enzyme that repairs the lesion the toxin creates. In this work, we showed that YwqJ is a cytidine deaminase that generates uracil in single-stranded DNA and mediates T7SSb-dependent antagonism, whereas EndoV^Bs^ is an endonuclease V that prefers uracil over hypoxanthine, processes genomic uracil in cells, and limits the damage YwqJ inflicts. To our knowledge, this is the first example of an enzyme encoded as a separate gene within a toxin-immunity operon that functions to repair, rather than neutralize, the damage inflicted by its associated effector. The structural basis of that preference remains unknown. Its consequences, however, are impactful in the ecology of *B. subtillis*, affecting how it interacts with other cells through the delivery of deaminase toxins, and how it responds to uracil-containing phages infection.

Toxin-immunity operons are generally understood to encode a poison and a means of neutralizing it, and in most characterized systems, a cognate immunity protein binds the toxin and prevents the lesion from forming(6–8). Enzymatic reversal of damage is an established alternative in interbacterial antagonisms. Tri1, the immunity determinant for the ADP-ribosyltransferase toxin Tre1 translocated by the T6SS, both occludes the toxin active site and removes ADP-ribose from the modified protein(37), and the antitoxin DarG likewise combines toxin binding with removal of ADP-ribose from DNA modified by DarT(38). In both cases, however, the enzymatic activity resides within the cognate immunity determinant itself. Immunity mechanisms are, in any case, more varied than occlusion alone. The immunity protein for Cpe1, an interbacterial cysteine protease that cleaves the ATPase domains of the type II topoisomerases GyrB and ParE, neutralizes the toxin by competing for its substrate rather than by blocking its active site(39). The hydrolase domain of Tri1 resembles DraG, an ADP-ribosylhydrolase with a housekeeping function unconnected to competition, and *tri1*-type immunity genes appear to have arisen from housekeeping genes by duplication on several independent occasions. Endonuclease V is likewise a repair enzyme of housekeeping origin, here resident in an antagonism locus.

Why *B. subtilis* requires a EndoV-dependent route to uracil is not immediately apparent, as the organism encodes a uracil-DNA glycosylase (Ung). The two enzymes differ in the intermediate they produce. Ung excises the base, leaving an abasic site that must be incised by an AP endonuclease before repair synthesis can begin. Abasic sites are also labile, and their spontaneous breakdown yields a strand break bearing a 3′ terminus that cannot prime synthesis. EndoV^Bs instead nicks the backbone 3′ of the lesion, generating a 3′-hydroxyl that DNA polymerase I can extend directly(24). Only the glycosylase route requires a second enzyme to act before synthesis begins. This distinction should matter when uracil accumulates faster than these steps can process it. Indeed, uracil-triggered chromosome fragmentation in *E. coli* requires Ung, whether uracil arises from dUTP misincorporation (40) or from a deaminase toxin (22). We propose that EndoV^Bs is suited to an acute uracil burden rather than to the sporadic deamination addressed by Ung (41).

Because EndoV^Bs^ acts on the lesion rather than on the toxin, its protection should be indifferent to which enzyme produced the uracil. The uracil processed in our heterologous system was in fact, generated by SsdA, a single-stranded DNA deaminase toxin unrelated to YwqJ (Fig. 3C). We speculate that EndoV^Bs^ affords *B. subtilis* a measure of protection against other DNA deaminase toxins delivered by competitors for which it carries no cognate immunity protein, a generic defense that a toxin-specific immunity protein cannot provide. Such a role would account for the occurrence of endonuclease V genes in genomes that lack *ywqJ* (Fig. 4C-E). Precedent for this exists in a different chemistry. The immunity protein Tri1 protects against ADP-ribosylating toxins by two mechanisms, active site occlusion and enzymatic removal of the modification, and only the latter confers protection against non-cognate effectors.

The restriction of PBS1 by EndoV^Bs^ bridges two systems typically regarded as functionally distinct: interbacterial antagonism, mediated by polymorphic toxins and immunity proteins, and antiviral defense, canonically carried out by dedicated antiphage systems in defense islands. Cross-talk between these arenas has been documented, but primarily through shared transcriptional regulation or surface trade-offs. In *Pseudomonas*, coordinated stress-sensing pathways coregulate both antibacterial effectors and antiphage defenses in response to envelope damage. In *Salmonella*, resistance to phages can lead to O-antigen loss that sensitizes cells to contact-dependent T6SS attack, creating an evolutionary trade-off between viral and bacterial defense.

In contrast, the association between phage defense and bacterial antagonism we describe requires no regulator to assemble two sets of defenses, and it exacts no trade-off. Loss of EndoV^Bs^ alters the outcome of infection by a phage that shares the lesion created by YwqJ. Adaptation to the toxin of a competitor thus carries consequences reaching beyond competition, and a locus assembled for antagonism proves to bear on phage susceptibility. This is also why we do not regard EndoV^Bs^ as a dedicated phage defense system. Such systems built on DNA glycosylases enriched within defense loci do exist. Brig1 excises α-glucosyl-hydroxymethylcytosine from the genome of phage T4, and the Dag families act on phages carrying modified guanine or thymidine. Those glycosylases also generate abasic sites in order to destroy phage DNA, whereas EndoV^Bs^ processes uracil without generating one. Consistent with this reading, the *endoV*^Bs^ locus lies outside any defense island, the protection it affords is smaller in magnitude than that conferred by dedicated systems, and its activation is independent of exposure to phages.

The co-occurrence of a DNA repair endonuclease within a polymorphic toxin locus reveals that toxin–immunity operons can encode repair to damage originated from sources independent of intoxication. Unlike regulatory co-activation (e.g., GRP/GUARD in *Pseudomonas*) or surface receptor trade-offs (*Salmonella*), this cross-protection operates through direct biochemical convergence on a common base lesion. Because EndoV recognizes the damage rather than the source, selection pressures, such as uracil-containing phages, can shape the maintenance and horizontal transfer of interbacterial weapon systems. We anticipate that bioinformatic and functional exploration of uncharacterized genes within polymorphic toxin operons will reveal that lesion-directed repair represents a widespread strategy for buffering bacteria against competitive and viral threats in polymicrobial ecosystems.

## Materials and Methods

### Bacterial strains and culture conditions

Unless otherwise stated, all cultures were grown in Lysogeny Broth (LB) liquid media or on solid media composed of LB solidified with 1.5% (w/v) agar. As required, media was supplemented with the following antibiotics for selection: carbenicillin (150 µg/mL), chloramphenicol (15 µg/mL), or kanamycin (50 µg/mL). *E. coli.* strain DH5α was used for plasmid cloning and maintenance. *E. coli.* strain BL21(DE3) was used for protein expression and intoxication assays. *Bacillus subtilis* strain JH642 was used for experiments involving *Bacillus*.

### Cloning, DNA assembly

All primers used in this study are listed [somewhere]. Restriction enzymes and Q5 2x PCR mastermix were obtained from New England Biolabs (NEB). Homology cloning was accomplished using the ClonExpress Ultra One Step Cloning Kit V2 from Vazyme. To co-express YwqJ_tox_ and YwqK, a pETDuet-1 expression construct was created by amplifying the toxin gene fragment and immunity gene from the *B. subtilis* genome.

### Protein purification

To purify his-tagged YwqJ_tox_ complexed with YwqK, 2 L of LB broth were inoculated with an overnight culture of *E. coli.* BL21(DE3) (pETDuet-1:: YwqJ_tox_ + YwqK) using a 1:100 dilution. The culture was grown to an OD_600_ of approximately 0.6, then induced with 1mM IPTG. Protein expression was then allowed to proceed by incubating the culture at 30°C for 4 hours with shaking. The culture was then pelleted by centrifugation at 4000 xG for 30 minutes. The pellet was then resuspended in 40 mL of lysis buffer (50 mM Tris-HCl pH 7.5, 400 mM NaCl, 5% (v/v) glycerol, 1mM DTT, 1 mg/mL lysozyme). The resuspended cell solution was then lysed at 4°C for 30 minutes with gentle agitation. The solution was then further lysed by sonication (five cycles of a 10 s pulse and 50 s rest). The lysate was clarified by centrifuging at 24,000 xG for 30 minutes at 4°C. Fast performance liquid chromatography (FPLC) was then used to purify the YwqJ_tox_-YwqK complex. Using an ÄKTA pure chromatography system (Cytivia), the clarified lysate was loaded onto a HisTrap HP affinity column (Cytivia). The column was washed with 20 column volumes (CV) of lysis buffer, followed by 20 CV of wash buffer (20 mM Tris-HCl pH 7.5, 350 mM NaCl, 1mM DTT). Bound YwqJ_tox_ - YwqK complex was then eluted using wash buffer with an imidazole gradient, increasing to 400 mM imidazole over 40 CV. All elution fractions were collected, and the fractions with eluted protein were pooled and retained.

To separate and purify YwqJ_tox_ from YwqK, a urea denaturing and refolding process was used. The pooled elution was then mixed with denaturing buffer (50 mM Tris-HCl pH 7.5, 500 mM NaCl, 8 M urea, 1mM DTT) to a volume of 50mL and incubated at 4°C for 16 hours with gentle agitation. A gravity flow column was packed with 1 mL Nuvia IMAC Ni-NTA resin (Bio-Rad) and equilibrated with denaturing buffer. The denatured protein was then passed through the column. The column was then washed with 10 CV renaturing buffer (20 mM Tris-HCl pH 7.5, 500 mM NaCl, 1mM DTT, 100nM ZnCl_2_) multiple times, with each 10 CV containing a decreasing amount of urea (8 M, 6 M, 4 M, 2 M, 1 M, 0 M) to allow on-column refolding of YwqJ_tox_. The column was then washed with 10 CV of wash buffer. YwqJ_tox_ was then eluted into multiple fractions, each using 1 mL of wash buffer with an increasing concentration of imidazole (50 mM, 100 mM, 200 mM, 300 mM, 400 mM). The protein concentration of each fraction was qualitatively determined using a Bradford assay (Bio-Rad), and the fractions with detectible protein were pooled. The pooled elution was then further purified with size-exclusion chromatography using a HiPrep 16/60 Sephacryl S-200 HR FPLC column in sizing buffer (20 mM Tris-HCl pH 7.5, 350 mM NaCl, 5% (v/v) glycerol, 1mM DTT). Elution fractions were collected, and the fractions with detectable protein within the elution volume range for a protein with the mass of YwqJ_tox_ were pooled. Purified YwqJ_tox_ was then distributed into multiple aliquots, flash frozen with liquid nitrogen, and stored at −80°C. Protein integrity and purity were assessed with SDS-PAGE and Spyro tangerine protein stain (Thermo Fisher).

To purify his-tagged EndoV^Bs^, an overnight culture of *E. coli.* BL21(DE3) (pET28b::ywqL) was diluted 1:100 in 2 L lB and grown to an OD_600_ of approximately 0.6. Protein expression and FPLC Ni-NTA affinity purification were performed as described above. After Ni-NTA purification, the pooled elation was concentrated on an Amicon Ultra-15 centrifugal filter unit with a 10 kDa molecular weight cutoff (MilliporeSigma). The sample was concentrated three times to exchange the elution buffer with storage buffer (20 mM Tris-HCl pH 7.5, 350 mM NaCl, 0.05% (v/v) Tween-20, 1mM DTT). Aliquots of purified EndoV^Bs^ were made, then flash frozen in liquid nitrogen and stored at −80°C. Protein integrity and purity were assessed with SDS-PAGE and Spyro tangerine protein stain (Thermo Fisher). EndoV^Bs^(D46A), EndoV^Bs^(4A), and *E. coli.* EndoV proteins were purified using the same method as EndoV^Bs^.

### DNA deamination assays

DNA oligonucleotide substrates containing a 5’ TET fluorophore were obtained from IDT. dsDNA substrates were prepared by annealing a reverse complement oligo to the fluorescently labeled substrate. Substrates were then purified using non-denaturing polyacrylamide gel electrophoresis. Biochemical reactions were performed in 10 µL of deamination buffer containing 20 mM Tris-HCl pH 7.5, 200 mM NaCl, 1 mM DTT, 8% (w/v) Ficoll-70. DNA substrate and YwqJ_tox_ were added in the concentration specified by Fig. X. The deamination reactions were incubated at 37°C for 1 hour. Next, 5 µL of UDG solution, consisting of UDG enzyme (New England Biolabs) diluted to 0.02 U/µL in 1x UDG buffer supplied by the manufacturer, was then added to each of the reactions and incubated at 37°C for an additional 30 minutes. To cleave the abasic sites generated by UDG, NaOH was added to a concentration of 100 mM followed by incubation at 95°C for 5 minutes. An equal volume of 2x TBE-Urea loading buffer (89mM Tris, 89 mM Boric acid, 2 mM EDTA, 12% Ficoll-70, 8M Urea, 0.3% Orange G) was mixed into each of the reactions. Reactions were incubated at 80°C for 5 minutes to fully denature the DNA substrates. Reaction products were loaded on a 20% acrylamide TBE-Urea gel and electrophoresed at 200V until the orange dye reached the bottom of the gel (30 - 45 minutes). The 5’ Tet labeled product was then visualized using a Li-Cor imager.

### EndoV DNA cleavage assays

DNA oligonucleotide substrates were acquired and prepared as previously described. EndoV protein and DNA substrate (type and concentration specified in Fig. X.) were combined in a final volume of 20uL with Endonuclease V assay buffer (5 mM Tris-HCl pH 7.5, 5 mM MgCl_2_, 5 mM DTT) and incubated at 37°C for 1 hour. The reactions were then mixed with 20 µL of 2x TBE-Urea loading buffer and incubated at 80°C for 5 minutes. The reaction products were then electrophoresed and visualized as described above.

### Flow cytometry

To track intoxication kinetics the proportion of GFP^+^ recipient cells in co-culture with wild-type donor cells was determined by flow cytometry. Cells were fixed in 2.5% paraformaldehyde for 15 minutes on ice and washed once in PBS. 100,000 cells per sample were analyzed on an LSR Fortessa X-20 flow cytometer using the FITC/GFP laser. Percent GFP^+^ cells was estimated by quantifying the higher fluorescing population in biphasic distributions.

### Fluorescence microscopy

To determine DNA content in individual cells, strains were fixed in 2.5% paraformaldehyde for 15 minutes on ice and washed once in TE. Cells were then stained with 1 µg/ml DAPI (4′,6-diamidino-2-phenylindole) in TE for 30 min on ice. Cells were spread onto agarose-coated slides to prevent movement and imaged on a Zeiss AxioImager Z1 equipped with a Hamamatsu EM-CCD camera.

DAPI fluorescence in 500-1000 cells per sample was measured using the open-source image processing package, Fiji. Background fluorescence was subtracted to give mean cellular DAPI fluorescence.

## Acknowledgements

The MHdM group was supported by a research grant from the National Science Foundation (Award Number: 2443063) and from the Welch Foundation (Award Number: C-2096-20220331). The BD group was supported by the National Institutes of Health (R01-GM135368)

We thank Dr. Cameron Glasscock for his thoughtful insights on the manuscript.

## Supplemental figures

**Supplementary Figure 1.**
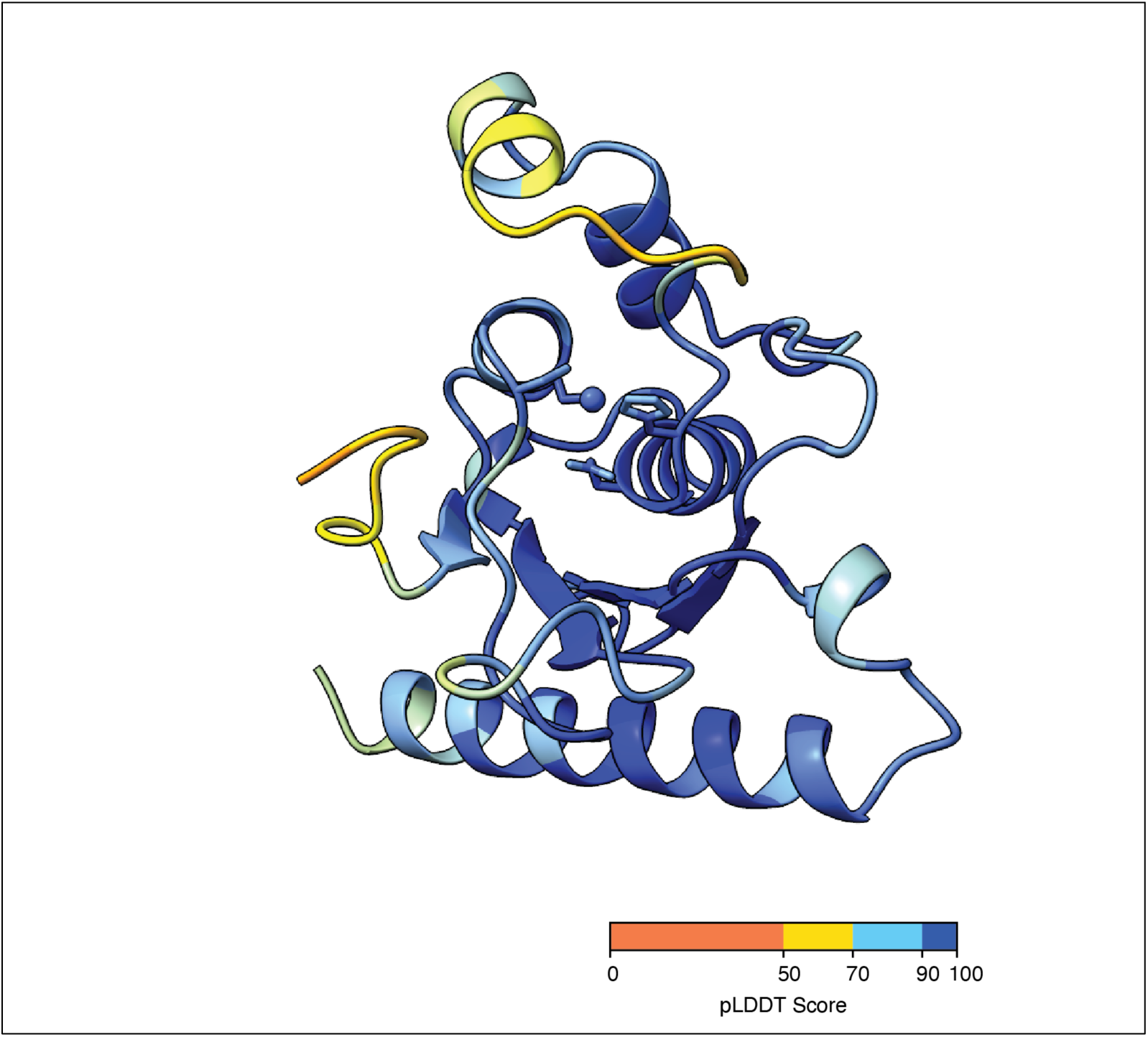
YwqJ toxin domain is predicted with high confidence. Structure prediction of YwqJ toxin domain by Alphafold3, with its residues colored by the corresponding pLDDT (predicted Local Distance Difference Test) score.

**Supplementary Figure 2:**
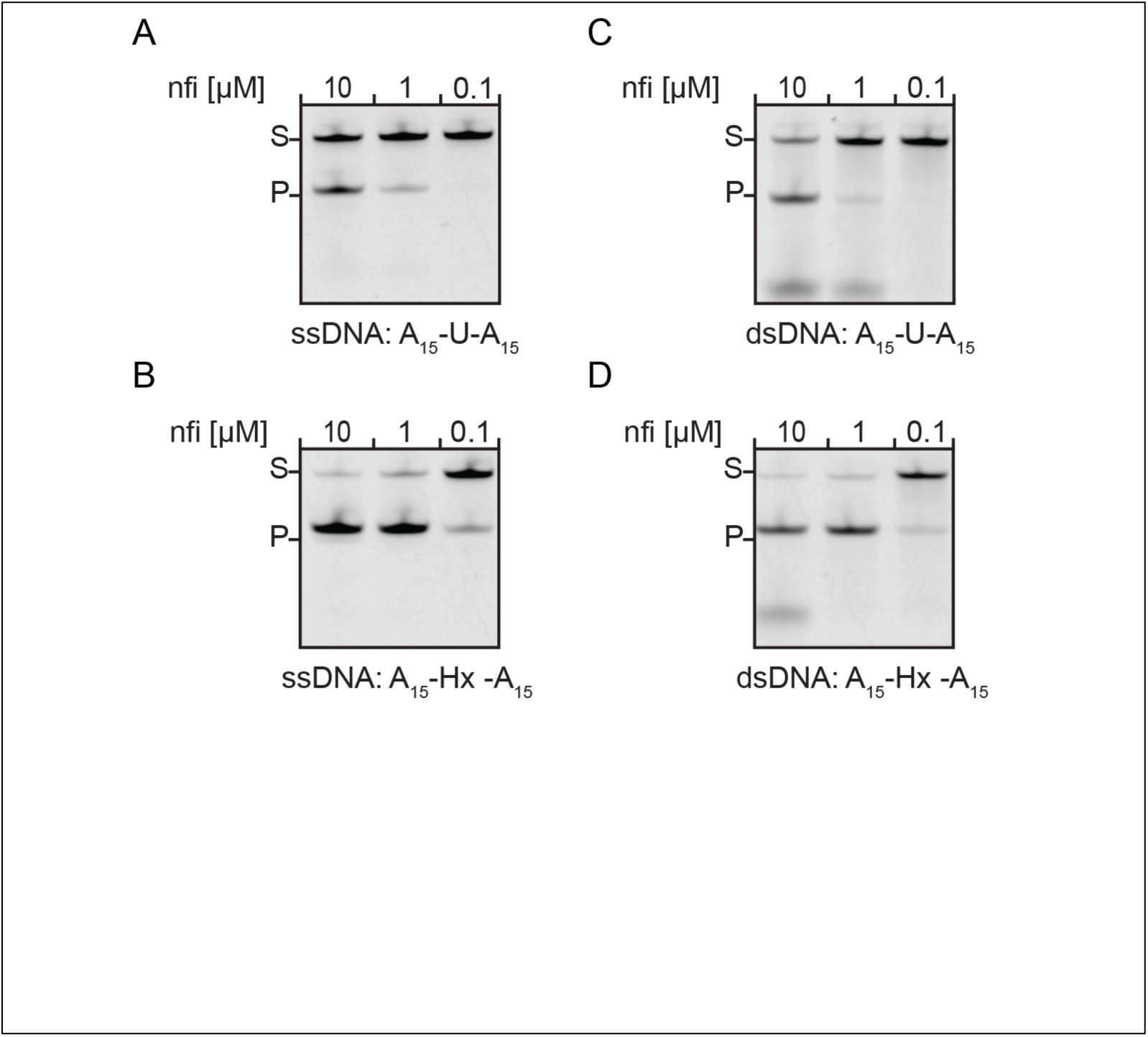
Nfi cleaves hypoxanthine-containing DNA in preference to uracil-containing DNA. Cleavage of 5′-TET-labeled oligonucleotide substrates by purified *E. coli* Nfi at the indicated enzyme concentrations, resolved by TBE-urea PAGE. **A**, single-stranded A_15_ - U-A_15_. **B**, single-stranded A_15_-Hx-A_15_. **C**, double-stranded A_15_-U-A_15_. **D**, double-stranded A_15_-Hx-A_15_. S, substrate; P, product. Nfi generated product from the hypoxanthine substrates at 1 and 0.1 μM, whereas cleavage of the uracil substrates was detected only at 10 μM. This is the inverse of the preference exhibited by EndoV^Bs^ (Fig. 3A, B).

**Supplementary Figure 3:**
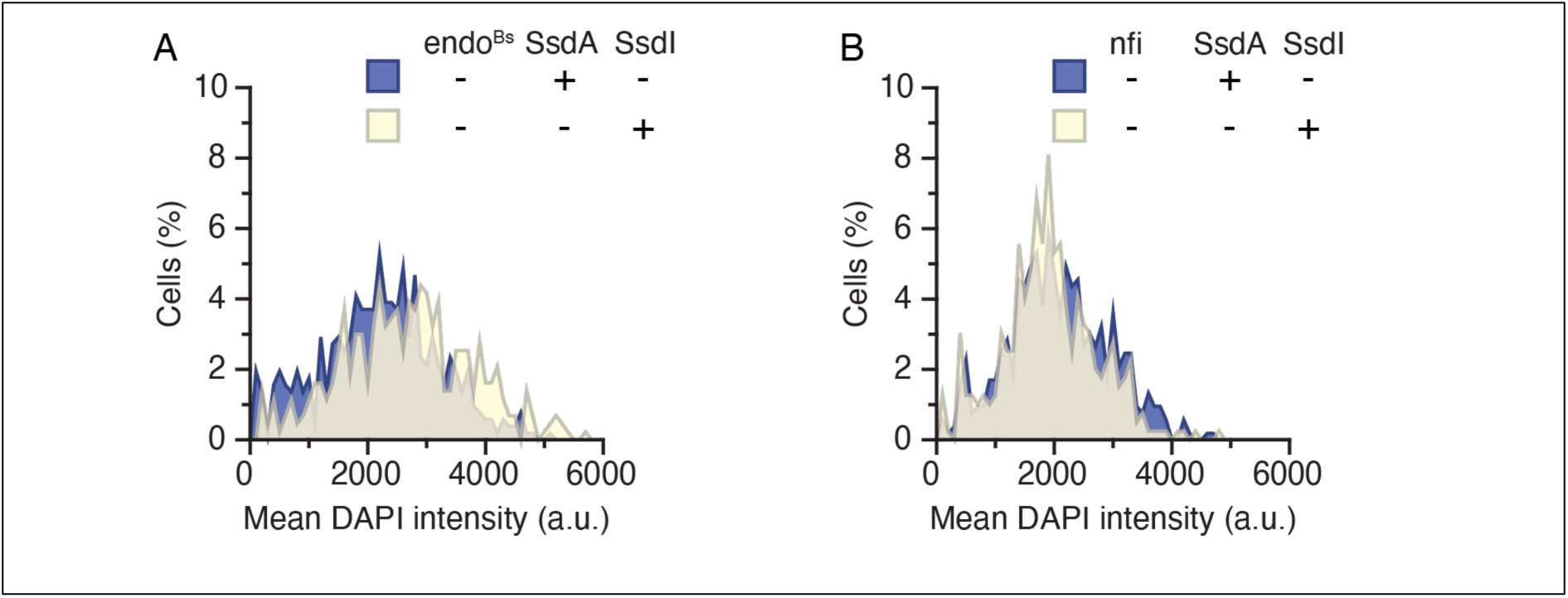
Distribution of nucleoid signal in the absence of endonuclease induction. Mean DAPI intensity per cell in *E. coli* Δ*ung* Δ*nfi* expressing active SsdA (blue) or SsdA neutralized by its cognate immunity protein SsdI (cream), with the endonuclease uninduced. **A**, cells carrying *endoV*^Bs^. **B**, cells carrying *nfi*. Values are background-subtracted per-cell means, binned and plotted as the percentage of cells per bin.

**Supplementary Figure 4:**
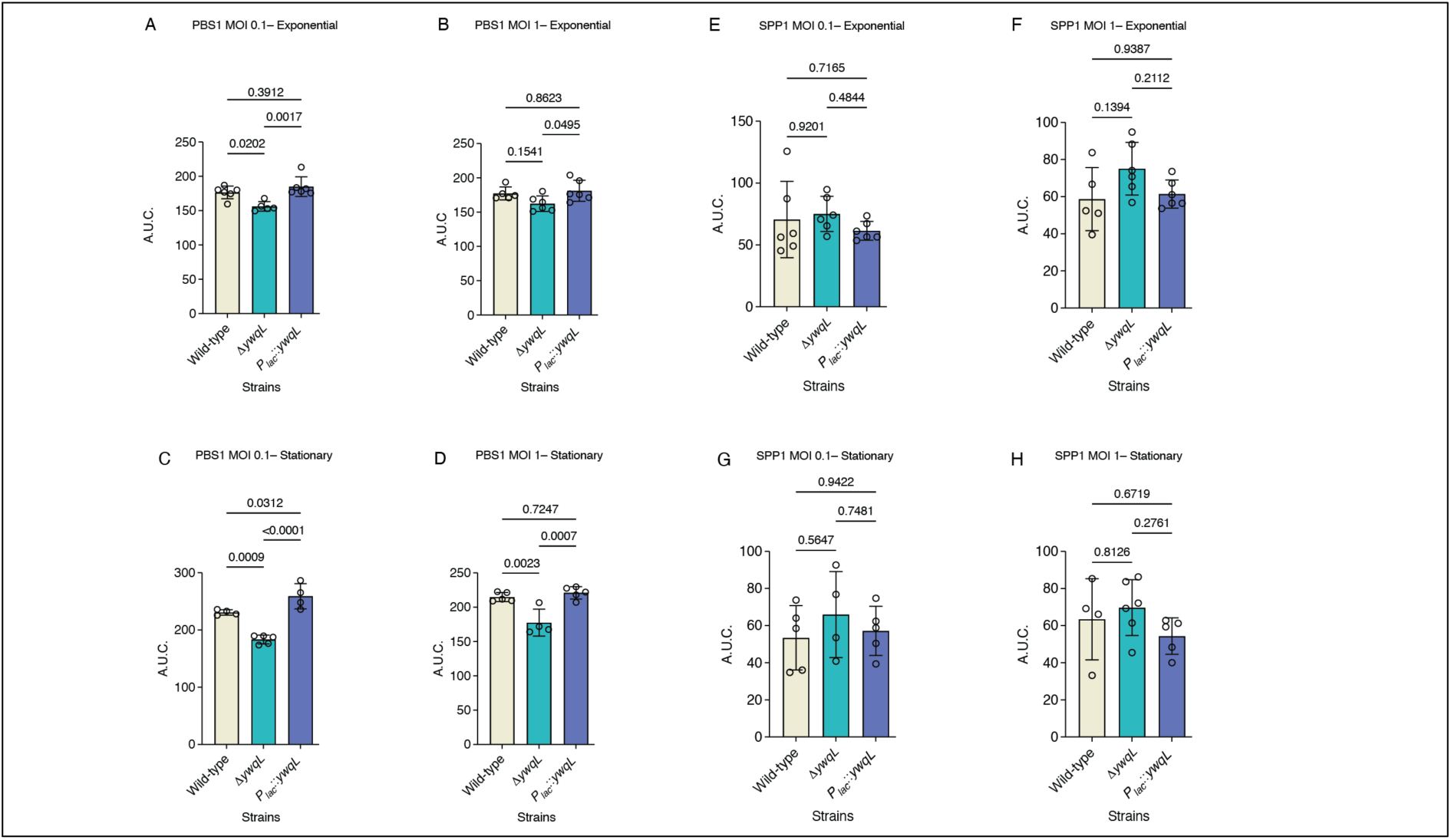
Area under the growth curve for PBS1- and SPP1-infected cultures. Area under the curve (A.U.C.) calculated from the growth curves shown in Fig. 5, for wild-type, Δ*ywqL* and P*_lac_*::*ywqL* strains. **A-D**, PBS1 at a multiplicity of infection of 0.1 (A, C) or 1 (B, D), with infection initiated in exponential (A, B) or stationary (C, D) phase. **E-H**, SPP1 under the same four conditions. Bars show the mean with individual replicates overlaid; error bars, s.d.; n = 4-6 biological replicates. Statistical significance was assessed by ordinary one-way ANOVA with Tukey’s multiple comparisons test.

